# Using genetics to examine a general liability to childhood psychopathology

**DOI:** 10.1101/409540

**Authors:** Lucy Riglin, Ajay K Thapar, Beate Leppert, Joanna Martin, Alexander Richards, Richard Anney, George Davey Smith, Kate Tilling, Evie Stergiakouli, Benjamin B. Lahey, Michael C O’Donovan, Stephan Collishaw, Anita Thapar

## Abstract

Psychiatric disorders show phenotypic as well as genetic overlaps. There are however also marked developmental changes throughout childhood. We investigated the extent to which, for a full range of early childhood psychopathology, a general “p” factor was explained by genetic liability, as indexed by multiple different psychiatric polygenic risk scores (PRS) and whether these relationships altered with age. The sample was a UK, prospective, population-based cohort with psychopathology data at age 7 (N=8161) and age 13 (N=7017). PRS were generated from large published genome-wide association studies. At both ages, we found evidence for a childhood “p” factor as well as for specific factors. Schizophrenia and attention-deficit/hyperactivity disorder (ADHD) PRS were associated with this general “p” factor at both ages but depression and autism spectrum disorder (ASD) PRS were not. Schizophrenia, ADHD and depression PRS were also associated with specific factors but there was evidence for developmental changes.

**Funding:** This work was supported by the Wellcome Trust (204895/Z/16/Z).

## Introduction

Most psychiatric disorders originate early in development (Rutter et al. 2006). Family and twin studies show that these disorders are heritable and many share genetic risk factors across diagnostic categories. For example, registry-based family and twin studies suggest that relatives of those with schizophrenia, attention-deficit/hyperactivity disorder (ADHD), autism spectrum disorder (ASD) and depression are at elevated risk for a broad range of psychopathology-not just for the same disorder as the affected probands (Doherty and Owen 2014). Recent molecular genetic studies also provide evidence of common variant genetic overlaps between different psychiatric disorders; for example genetic correlations across schizophrenia, ADHD, ASD and mood disorders have been shown to range from 0.30 to 0.93 (Schork et al. 2019). Current psychiatric nosology does not take into account these complex patterns of shared inheritance across phenotypes (Doherty and Owen 2014).

While the genetic architecture of psychiatric disorders is complex, indicators of (common variant) genetic liability for psychiatric disorders including schizophrenia, ADHD, ASD and depression can be measured in individuals using polygenic risk scores (PRS) (Sullivan et al. 2018). Studies using this method also support the existence of shared genetic risks. For example, schizophrenia PRS are associated with increased liability to other disorders (including ADHD, ASD, depression and anxiety) and broader categories of psychiatric phenotypes (e.g. neurodevelopmental traits in childhood) (Riglin et al. 2017; Schizophrenia Working Group of the Psychiatric Genomics Consortium 2014). Although it is now widely accepted that psychiatric disorder risk alleles are shared across disorders, how such risks have a shared impact on psychopathology remains unclear.

While different forms of psychopathology are considered as discrete categories for clinical purposes, it has long been known that different disorders show strong phenotypic (as well as genetic) overlap. Factor analyses of child and adult psychopathology have observed that these phenotypic overlaps can be largely explained by a single, latent general factor that reflects general liability to psychopathology, known as the “p” factor (Lahey et al. 2011; Caspi and Moffitt 2018). In children, factor analyses show that different forms of psychopathology converge onto specific “emotional” (internalizing) and “behavioural” (externalizing) factors as well as a general “p” factor. However, such studies typically have not included measures of ASD and do not examine whether associations change with age (Ronald 2019). In adulthood, recent studies have highlighted the presence of a specific “thought disorder” factor, as well as “emotional” and “behavioural” factors that aggregate into one general psychopathology dimension (Caspi and Moffitt 2018). Indeed, it has been proposed recently that the general liability factor “p” could account for the relative lack of clinical and treatment specificity and shared risk factors across different forms of psychopathology (Caspi and Moffitt 2018). However, this requires further empirical testing because there are some unique aspects to different types of psychopathology. For example, stimulant medication is effective for ADHD but not for anxiety disorder or ASD suggesting that there are also important biological differences between disorders.

In this study, we set out to examine associations between risk alleles for several different major psychiatric disorders, indexed by PRS, and the ‘general psychopathology’ factor and whether these relationships were altered as children became older. Unlike previous studies of psychopathology, we included ASD problems and hypothesised that we would identify a “neurodevelopmental” factor as well as “emotional” and “behavioural” factors. We investigated PRS for schizophrenia, ADHD, ASD and depression, derived from large, publicly available genome-wide association studies, because these PRS have previously been shown to be associated with childhood as well as adult psychopathology (Poletti and Raballo 2018; St Pourcain et al. 2018; Brikell et al. 2018; Rice et al. 2019). First, we hypothesized that in childhood, all of these psychiatric genetic risk scores would be associated with the general “p” factor. Our second hypothesis was that PRS would show additional phenotype-specific associations; specifically, that we would observe associations between ASD and ADHD PRS with the specific “neurodevelopmental” factor and depression PRS with the specific “emotional” factor.

## Methods

### Sample

The Avon Longitudinal Study of Parents and Children (ALSPAC) is a well-established prospective, longitudinal birth cohort study. The enrolled core sample consisted of 14,541 mothers living in Avon, England, who had expected delivery dates of between 1^st^ April 1991 and 31^st^ December 1992. Of these pregnancies 13,988 children were alive at 1 year. When the oldest children were approximately 7 years of age, the sample was augmented with eligible cases who had not joined the study originally, resulting in enrolment of 713 additional children. The resulting total sample size of children alive at 1 year was N=14,701. Genotype data were available for 8,365 children following quality control. Ethical approval for the study was obtained from the ALSPAC Ethics and Law Committee and the Local Research Ethics Committees. Full details of the study, measures and sample can be found elsewhere (Boyd et al. 2013; Fraser et al. 2013). Please note that the study website contains details of all the data that is available through a fully searchable data dictionary (http://www.bris.ac.uk/alspac/researchers/data-access/data-dictionary). Where families included multiple births, we included the oldest sibling. We used childhood measures at age 7 years and age 13 years.

### Polygenic risk scores

Polygenic risk scores (PRS) were generated as the weighted mean number of disorder risk alleles in approximate linkage equilibrium, derived from imputed autosomal SNPs using PRSice (Euesden et al. 2015). Scores were standardized using Z-score transformation. Risk alleles were defined as those associated with case-status in recent large consortia analyses of schizophrenia (40,675 cases and 64,643 controls) (Pardinas et al. 2018), ADHD (19,099 cases and 34,194 controls) (Demontis et al. 2019), ASD (18,381 cases and 27,969 controls) (Grove et al. 2019) and depression (135,458 cases and 344,901 controls) (Wray et al. 2018). In the primary analyses we defined risk alleles as those associated at p<0.05 as this threshold has previously been shown to maximally capture phenotypic variance for schizophrenia (Schizophrenia Working Group of the Psychiatric Genomics Consortium 2014); associations across a range of p-thresholds are shown in Supplementary Figure 1 in the online data supplement. Genotyping details as well as full methods for generating the PRS are presented in the Supplementary Material. PRS were available for 68% of individuals (N=5518/8161) who also had phenotypic data at age 7. Sensitivity analyses were conducted using inverse probability weighting (Seaman and White 2013) to assess the impact of missing genetic data (see Supplementary Material).

**Figure 1.**
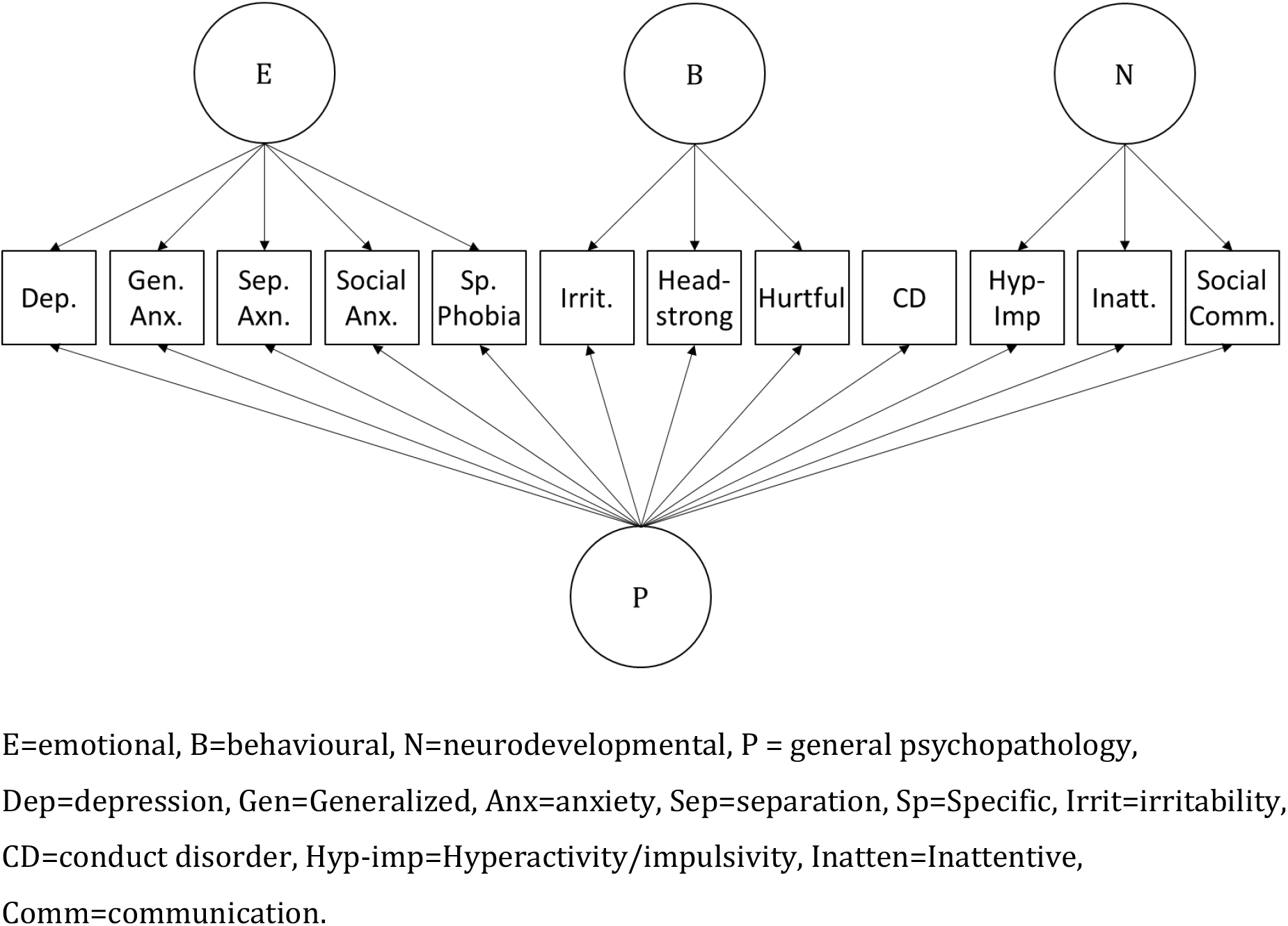
Bifactor model

### Outcomes

Psychopathology was assessed using parent reports at the approximate age of 7/8 years old and age 13. Emotional problems were assessed using the Development and Well-Being Assessment (DAWBA) (Goodman et al. 2000) (individual item range 0-2) for depression (12 items), generalized anxiety (7 items), separation anxiety (10 items), social anxiety (6 items) and specific phobia (7 items). Behavioural problems were assessed using the DAWBA for conduct disorder (7 items) and the three components of oppositional defiant disorder: irritability (3 items), headstrong (4 items) and hurtful (2 items) behaviours. Neurodevelopmental problems were assessed using the DAWBA for activity/impulsivity (9 items) and inattention (9 items) for ADHD problems. Additionally, the Social and Communication Disorders Checklist (Skuse et al. 2005) was used for social-communication problems related to ASD (12 items; individual item range 0-2). Prorated scores (the mean score multiplied by total possible number of items in the scale) were calculated for individuals with <30% missingness. Descriptive statistics and correlations between variables are given in Supplementary Tables I and II.

### Analysis

In-line with previous factor analysis work, we first used confirmatory factor analysis to fit bifactor models (including both general and specific factors, with correlations between factors fixed to zero) at each age. Model fit was assessed using a variety of indices including the comparative fit index (CFI, >0.95 considered good fit), Tucker-Lewis index (TFI, >0.95 considered good fit) and the root-mean-square error of approximation (RMSEA, <0.06 considered good fit) (Hu and Bentler 1999). Comparisons of model fit with different models are available from the first author. Omega reliability coefficients were calculated to assess model reliability (Rodriguez et al. 2016).

Associations with PRS were examined in two steps. First, factors were fixed based on the best fitting model before associations with PRS were examined (i.e. PRS did not affect the factor structure). Second, schizophrenia, ADHD, ASD and depression PRS were included simultaneously (as were all factors within a given model). Associations with PRS were therefore investigated in one test at age 7 years and one test at age 13 years. Sensitivity analyses were conducted to investigate associations for each of the four PRS separately (univariable analyses). Analyses were conducted in Mplus using a maximum likelihood parameter estimator for which standard errors are robust to non-normality (MLR) (Muthén and Muthén 1998-2012).

## Results

### Factor model

At age 7 a bifactor model including a general psychopathology factor “p” as well as three ‘specific’ “emotional”, “behavioural” and “neurodevelopmental” factors – shown in Figure 1 and Table 1 - fit the data well (CFI=0.973, TFI=0.959, RMSEA=0.038). Omega reliability coefficients, also shown in Table 1, indicated that the inclusion of the specific factors explained additional variance beyond that explained by the “p” factor (ω; compared to the general factor alone ω_H_). These coefficients indicated that for ADHD and ASD problems, most of the variance was explained by the general “p” factor. In contrast, for anxiety/mood problems, most of the variance was explained by the specific “emotional” factor (variance explained was similar regardless of whether or not the general factor was partialled out: ω_HS_ compared to ω_S_). Finally, for oppositional defiant problems, both the general “p” factor and specific “behavioural” factor captured variance.

**Table I.** Factor loadings and omega reliability coefficients at age 7

|  | P | E | B | N |
| --- | --- | --- | --- | --- |
| Factor loadings |  |  |  |  |
| Depression | 0.292 | 0.311 |  |  |
| Generalized anxiety | 0.230 | 0.566 |  |  |
| Separation Anxiety | 0.296 | 0.500 |  |  |
| Social anxiety | 0.206 | 0.402 |  |  |
| Specific phobia | 0.158 | 0.447 |  |  |
| Irritability | 0.731 |  | 0.416 |  |
| Headstrong | 0.824 |  | 0.374 |  |
| Hurtful | 0.622 |  | 0.351 |  |
| Conduct disorder | 0.557 |  |  |  |
| Hyperactivity-impulsivity | 0.697 |  |  | 0.535 |
| Inattention | 0.608 |  |  | 0.564 |
| Social-communication | 0.795 |  |  | 0.199 |
| Omega reliability coefficients |  |  |  |  |
| Omega ($\omega$) | 0.944 | | | |
| Omega hierarchical ($\omega_H$) | 0.554 | | | |
| Omega subscale ($\omega_S$) | | 0.933 | 0.947 | 0.894 |
| Omega hierarchical subscale ($\omega_{HS}$) | | 0.897 | 0.420 | 0.054 |

P = general psychopathology, E=emotional, B=behavioural, N=neurodevelopmental

Female sex was negatively associated with the general psychopathology “p” factor (β=-0.106, SE=0.014, p<0.001), positively associated with the “emotional” factor (β=0.117, SE=0.017, p<0.001), not strongly associated with the “behavioural” factor (β =0.028, SE=0.024, p=0.231) and negatively associated with the “neurodevelopmental” factor (β =-0.147, SE=0.018, p<0.001). At age 13 a similar pattern of results was found apart from a lower reliability coefficient (Omega hierarchical (ωH)) for the general factor and that the behavioural and ADHD/ASD factors explaining a larger proportion of the variance in scores in these domains than at age 7 (see Supplementary Table III).

### Genetic risk

Results of multivariable PRS association with the psychopathology factors are shown in Table 2. At both age 7 schizophrenia and ADHD PRS were associated with the general psychopathology “p” factor, while ASD and depression PRS were not. Schizophrenia PRS were also associated with the specific “emotional” factor; depression PRS were weakly associated with the “emotional” factor and ADHD PRS were negatively associated with the specific “emotional” factor. There was no strong evidence of associations between PRS and the behavioural or neurodevelopmental factor.

**Table II.** Multivariable associations between genetic risk and the factor model at ages 7 and 13 years

| Age 7 | Schizophrenia PRS |  |  | ADHD PRS |  |  | ASD PRS |  |  | Depression PRS |  |  | Total R<sup>2</sup> |
| --- | --- | --- | --- | --- | --- | --- | --- | --- | --- | --- | --- | --- | --- |
| | $\beta$ | SE | p | $\beta$ | SE | p | $\beta$ | SE | p | $\beta$ | SE | p | |
| General psychopathology | 0.042 | 0.018 | 0.016 | 0.087 | 0.019 | <0.001 | 0.002 | 0.018 | 0.912 | 0.026 | 0.017 | 0.144 | 0.011 |
| Emotional problems | 0.040 | 0.019 | 0.035 | -0.046 | 0.020 | 0.021 | 0.007 | 0.019 | 0.713 | 0.035 | 0.019 | 0.073 | 0.005 |
| Behavioural problems | 0.015 | 0.028 | 0.577 | -0.050 | 0.034 | 0.143 | -0.022 | 0.029 | 0.444 | -0.017 | 0.028 | 0.539 | 0.004 |
| Neurodevelopmental problems | -0.002 | 0.022 | 0.539 | 0.041 | 0.025 | 0.109 | -0.021 | 0.023 | 0.362 | -0.002 | 0.022 | 0.912 | 0.002 |
| Age 13 | Schizophrenia PRS |  |  | ADHD PRS |  |  | ASD PRS |  |  | Depression PRS |  |  | Total R<sup>2</sup> |
| | $\beta$ | SE | p | $\beta$ | SE | p | $\beta$ | SE | p | $\beta$ | SE | p | |
| General psychopathology | 0.056 | 0.023 | 0.015 | 0.095 | 0.020 | <0.001 | -0.016 | 0.019 | 0.398 | 0.012 | 0.020 | 0.559 | 0.013 |
| Emotional problems | -0.016 | 0.020 | 0.421 | -0.015 | 0.020 | 0.437 | 0.019 | 0.019 | 0.313 | 0.056 | 0.018 | 0.002 | 0.004 |
| Behavioural problems | -0.037 | 0.036 | 0.308 | -0.019 | 0.031 | 0.536 | 0.004 | 0.032 | 0.899 | -0.001 | 0.031 | 0.985 | 0.002 |
| Neurodevelopmental problems | 0.004 | 0.041 | 0.921 | 0.061 | 0.024 | 0.011 | -0.010 | 0.025 | 0.706 | -0.005 | 0.028 | 0.852 | 0.004 |

ADHD=attention-deficit/hyperactivity disorder, ASD=autism spectrum disorder, PRS=polygenic risk score.
Betas represent a 1 standard deviation increase in factor score per 1 standard deviation increase in PRS.

As at age 7, at age 13 schizophrenia and ADHD PRS were associated with the general psychopathology “p” factor, while ASD and depression PRS were not. Notable differences were a stronger association between the depression PRS and the emotional problems factor and between ADHD PRS and the neurodevelopmental problems factor and that schizophrenia and ADHD PRS were no longer associated with the emotional factor.

There was no strong evidence of associations between ASD PRS and any of the factors.

### Sensitivity analyses

Entering PRS in univariable (instead of multivariable) analyses revealed a similar pattern of results, with the exception that the association between depression PRS and general psychopathology was stronger when the other PRS were not included in the model (age 7 β=0.041, SE=0.017, p=0.019, see Supplementary Table IV for full results). Associations for schizophrenia, ASD and ADHD PRS were consistent regardless of whether or not the other PRS were controlled for.

Using inverse probability weighting to assess the impact of missing genetic data revealed a similar pattern of results (see Supplementary Material).

## Discussion

In this study we first set out to examine whether genetic liability as indexed by PRS for different psychiatric disorders would be associated with a general childhood psychopathology “p” factor that captures the main forms of psychopathology including ASD problems and also as to whether these associations varied with age. This has not previously been tested. As expected, given previous factor analysis studies, a bifactor model fit the data well. This included the general “p” factor and specific “emotional” and “behavioural” factors. Uniquely, we also included ASD symptoms and additionally observed a third “neurodevelopmental” factor that captured variance for ADHD and ASD problems although, most of the variance for these problems was captured by the general “p” factor.

Our first hypothesis was partially but not entirely supported: schizophrenia and ADHD PRS were associated with a general psychopathology “p” factor both at age 7 years and at age 13 years but associations were not observed for ASD and depression PRS at either age. This suggests that there may be a more complex explanation to the structure of psychopathology and its aetiology than a single unifying dimension.

Our second hypothesis was that PRS would show additional phenotype-specific associations; specifically, that we would observe associations between ASD and ADHD PRS with the specific “neurodevelopmental” factor and depression PRS with the specific “emotional” factor. This was also partially supported, with expected specific associations for ADHD and depression at age 13. In addition, at the younger age point (7 years) we found schizophrenia PRS to be associated with the specific “emotional” factor and ADHD PRS were negatively associated with the “emotional” factor.

For schizophrenia PRS several studies have now reported associations with psychopathology as well as IQ and social traits from early childhood onwards (Riglin et al. 2017; Poletti and Raballo 2018; Riglin et al. 2018). Our findings suggest that these associations may be driven mainly, but not exclusively by a general liability to psychopathology. Recent work in this sample that focused on psychosis-related and emotional problems, suggests the same to be true for associations between schizophrenia PRS and other forms of psychopathology (i.e. psychotic experiences, depression and anxiety) in later adolescence (Jones et al. 2018). This is consistent with previous suggestions that risk factors for rarer forms of psychopathology, such as schizophrenia, are also associated with (non-specific) risk for common psychopathology via a single general “p” factor (Lahey et al. 2017). However, we also observed association between schizophrenia PRS and the specific “emotional” factor, at age 7 suggesting some specificity in associations between genetic liability to schizophrenia and depression/anxiety, at least in early childhood.

Results for depression PRS also suggest that a single liability model for psychopathology is insufficient. Depression PRS were not independently associated with the general “p” factor but were associated with the specific “emotional” factor at age 13. This association was much stronger at this age than at age 7 years, suggesting that developmental change is likely important. This is consistent with previous work in another UK cohort that found depression PRS to be associated with emotional problems in adulthood but not in childhood (Riglin et al. 2018).

ADHD PRS was associated with both the general psychopathology “p” factor at both ages and also with the specific “neurodevelopmental” factor at age 13 - consistent with another population-based study that investigated ADHD PRS. That study of Swedish children aged 9-12 years observed associations between ADHD PRS and a latent general “p” factor encompassing emotional, behavioural and neurodevelopmental problems, which was also associated with specific ADHD hyperactive/impulsive problems – although they did not investigate a specific “neurodevelopmental” factor (Brikell et al. 2018). The findings taken together suggest that while ADHD genetic liability contributes to child psychopathology via a general liability or “p” factor, there is also additional specificity in its associations with ADHD/neurodevelopmental problems. Unexpectedly, we found a negative association between ADHD PRS and the specific “emotional” factor at age 7 – similar findings were observed for social phobia in previous work (Brikell et al. 2018) although it is not clear why. It may be that during early childhood (age 7) genetic liability for ADHD is so strongly captured by the general “p” factor that the overlap with emotional symptoms is explained by this general factor and the remaining loading becomes negative. Another possibility is that the genetic and clinical overlap between ADHD and emotional problems (depression, anxiety) changes after mid-late adolescence. Finally, for ASD PRS no associations with either the general or any specific factor were found at either age. Previous studies have found associations between ASD PRS and social-communication problems related to ASD (one of our indices of neurodevelopmental problems) (St Pourcain et al. 2018) but our findings suggest that ASD common variants (at least as far as can be estimated using currently available GWAS discovery sample sizes) do not contribute to a broader (latent) liability for “neurodevelopmental” problems or general childhood psychopathology. This is somewhat surprising given that family and twin studies, the latest ASD patient GWAS and studies of rare genetic mutations all suggest considerable genetic overlap between ASD and ADHD diagnoses (Rommelse et al. 2010). We speculate that this could in part be due to the relatively small ASD discovery GWAS and the typically weaker associations with trait measures than diagnoses. In addition, genetic findings for ASD PRS do not parallel observations for many other neuropsychiatric PRS (e.g. ADHD, schizophrenia). For example, unlike ADHD PRS, they predict higher IQ in population-based samples (Grove et al. 2019). We also found that as children got older the proportion of variance in symptom scores captured by specific (particularly the behavioural and ADHD/ASD) domains increased with a corresponding small decrease for symptom variance exclusively captured by the general factor. This needs further study but reinforces the need to take a developmental perspective.

Taken together our findings, using neuropsychiatric PRS, provide evidence in favour of a general liability to childhood psychopathology but also the importance of specific domains of psychopathology and the importance of taking development into account when examining these associations. For neurodevelopmental and early behavioural problems we observed much of the phenotypic variance was captured by a general psychopathology factor particularly during early childhood but with evidence for an increasing contribution for specific factors as children became older. However, for emotional problems, a specific factor accounted for the majority of the variance in these phenotypes at both ages but the association with the depression PRS was stronger as children got older.

Our findings should be considered in light of a number of limitations. First, the sample is a longitudinal birth cohort study that suffers from non-random attrition, whereby children with higher PRS and higher levels of psychopathology are more likely to drop out of the study (Taylor et al. 2018). Second, our models were statistically driven - although theoretically informed - and are therefore dependent on the properties of the measures that are included in the models; while our bifactor model was the best fitting model that we tested, this does not mean that this is the best possible model. We had to restrict our measures to parent-rated problems given the young age of the children - this is typical practice. Finally, it is also important to note that PRS currently only explain a small proportion of genetic liability to psychiatric disorders (Schizophrenia Working Group of the Psychiatric Genomics Consortium 2014) and that environmental risk factors and shared symptoms (as well as other non-common, non-additive genetic factors not captured by existing PRS) will also contribute to phenotypic overlaps (Lahey et al. 2011).

### Conclusion

Genetic liability for schizophrenia and ADHD as indexed by PRS are associated with a ‘general psychopathology’ factor and this association persists from early to mid-childhood age (age 7 to age 13 years). In addition, however, by mid-childhood, associations between ADHD PRS and a specific neurodevelopmental factor and depression PRS and a specific emotional factor emerge. Our conclusion from these findings is that, using genetic liability, a unidimensional perspective for psychopathology is helpful but insufficient and that there is also specificity for types of psychopathology that emerges throughout childhood and that a developmental perspective is valuable for investigating this.

## Acknowledgements

We acknowledge the members of the Psychiatric Genomics Consortium for the publicly available data used as the discovery samples in this article. We thank the research participants and employees of 23andMe, Inc. for their contribution to this study. We thank all the families who took part in this study, the midwives for their help in recruiting them, and the entire Avon Longitudinal Study of Parents and Children team, which includes interviewers, computer and laboratory technicians, clerical workers, research scientists, volunteers, managers, receptionists, and nurses. The UK Medical Research Council, grant 102215/2/13/2 from the Wellcome Trust, and the University of Bristol provide core support for the Avon Longitudinal Study of Parents and Children. Genome-wide association study data were generated by Sample Logistics and Genotyping Facilities at the Wellcome Trust Sanger Institute and Laboratory Corporation of America using support from 23andMe. This study was supported by the Wellcome Trust (204895/Z/16/Z). Dr Martin was supported by the Wellcome Trust (Grant No: 106047).

## Conflict of Interest Disclosures

None.

