## Supplementary Materials for "Using genetics to examine a general liability to childhood psychopathology"

**Supplementary Material**

### Generating polygenic risk scores

In total 9912 ALSPAC children were genotyped using the Illumina HumanHap500-quad genotyping array. Individuals were excluded on the basis of gender mismatches; minimal or excessive heterozygosity, disproportionate levels of individual missingness (>3%), insufficient sample replication (IBD <0.8), non-European ancestry (assessed by multidimensional scaling analysis and compared with Hapmap II) and cryptic relatedness (IBD > 0.1). SNPs were excluded based on minor allele frequency (<1%), call rate (<95%) or evidence for violations of Hardy-Weinberg equilibrium (P < 5E-7). Imputation was conducted by the ALSPAC team using Impute V2.2.2 against the 1000 genomes reference panel (Phase 1, Version 3: all polymorphic SNPs excluding singletons), using all 2186 reference haplotypes (including non-Europeans). SNPs were subsequently filtered based on minor allele frequency (<1%) and imputation quality (INFO<0.8). Following quality control and limiting individuals to one child per family, genetic data were available for N=7975.

Genome-wide association study (GWAS) summary statistics used to generate PRS were filtered to remove SNPs that were palindromic, insertions/deletions, non-autosomal, INFO score <0.8, missing in N>1 study and duplicates (https://github.com/ricanney). Depression results for 23andme (75,607 cases and 231,747 controls) (Hyde et al. 2016) and the other samples included in the latest depression GWAS (Wray et al. 2018) (PGC29, deCODE, Generation Scotland, GERA, iPSYCH, and UK Biobank) were meta-analysed in METAL. PRS were generated in ALSPAC using PRSice (Euesden et al. 2015); SNPs were clumped with an $R^2$ threshold of 0.1 and a distance threshold of 1000kb and excluding the extended major histocompatibility complex (MHC; chromosome 6: 26-33Mb) due to the high linkage disequilibrium (LD) within this region. Polygenic risk scores were standardized using Z-score transformation. Correlations between the different PRS are shown in Supplementary Table VI.

**Inverse probability weighting**

Inverse probability weighting (IPW) was used to assess the impact of missing genetic data and has been recommended over alternative methods such as multiple imputation in situations where whole blocks of data are missing for a large proportion of individuals (Seaman et al. 2012). Weights were derived from a logistic regression analysis of missing genetic data for those in the 'core' ALSPAC sample (N=6298/13793) for a set of measures assessed in pregnancy with minimal missingness: child gender (0% missing), child birth weight (1.3% missing: singly imputed) and maternal age (0% missing). Differences between those with and without genetic data for these variables are shown in Supplementary Table VII. Analyses conducted using IPW to address any potential bias caused by only a subsample having genetic data revealed a similar pattern of results (see Supplementary Table VIII).

**Supplementary Figure 1.** Multivariable associations between the bifactor model at age 7 years and polygenic risk scores of varying p-value thresholds

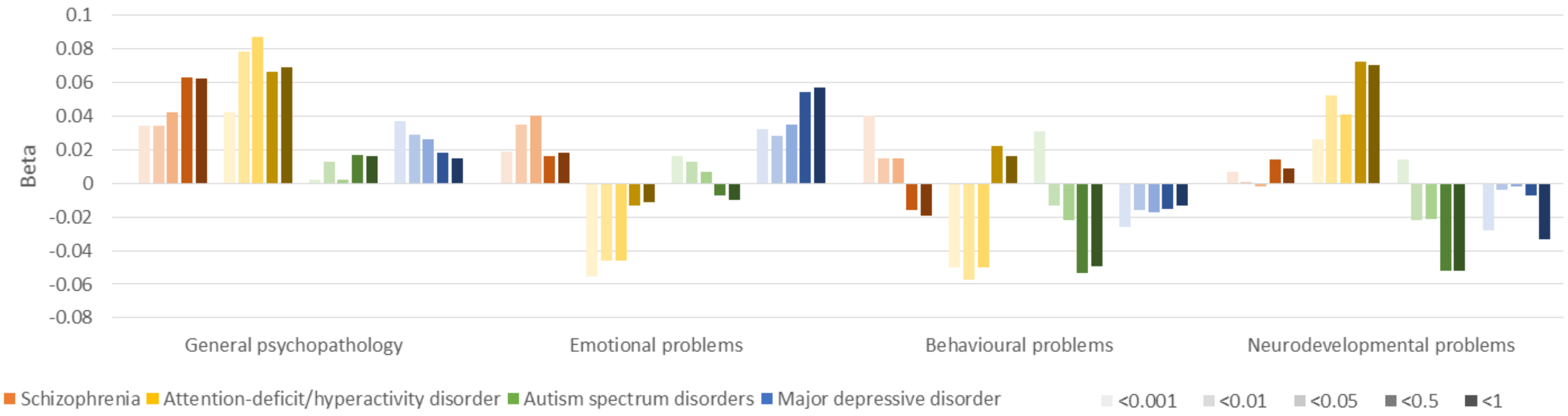

**Supplementary Table I.** Descriptive statistics including correlations between variables at age 7

| | 1. | 2. | 3. | 4. | 5. | 6. | 7. | 8. | 9. | 10. | 11. | 12. |
|---|---|---|---|---|---|---|---|---|---|---|---|---|
| 1. Depression | 1 | | | | | | | | | | | |
| 2. Generalized anxiety | 0.280 | 1 | | | | | | | | | | |
| 3. Separation Anxiety | 0.267 | 0.330 | 1 | | | | | | | | | |
| 4. Social anxiety | 0.165 | 0.281 | 0.250 | 1 | | | | | | | | |
| 5. Specific phobia | 0.111 | 0.288 | 0.297 | 0.229 | 1 | | | | | | | |
| 6. Irritability | 0.278 | 0.218 | 0.255 | 0.182 | 0.134 | 1 | | | | | | |
| 7. Headstrong | 0.219 | 0.170 | 0.237 | 0.145 | 0.111 | 0.758 | 1 | | | | | |
| 8. Hurtful | 0.198 | 0.153 | 0.201 | 0.154 | 0.105 | 0.602 | 0.644 | 1 | | | | |
| 9. Conduct disorder | 0.163 | 0.131 | 0.157 | 0.078 | 0.045 | 0.380 | 0.452 | 0.418 | 1 | | | |
| 10. Hyperactivity-impulsivity | 0.188 | 0.167 | 0.208 | 0.119 | 0.116 | 0.475 | 0.593 | 0.407 | 0.395 | 1 | | |
| 11. Inattention | 0.217 | 0.168 | 0.196 | 0.198 | 0.115 | 0.427 | 0.499 | 0.342 | 0.346 | 0.725 | 1 | |
| 12. Social-communication | 0.217 | 0.167 | 0.215 | 0.185 | 0.156 | 0.589 | 0.655 | 0.472 | 0.445 | 0.661 | 0.594 | 1 |
| Polygenic risk scores | | | | | | | | | | | | |
| Schizophrenia | -0.005 | 0.022 | 0.037 | 0.064 | 0.031 | 0.048 | 0.039 | 0.026 | 0.005 | 0.025 | 0.028 | 0.053 |
| Attention-deficit/hyperactivity disorder | 0.012 | 0.002 | 0.010 | -0.019 | -0.001 | 0.029 | 0.060 | 0.048 | 0.089 | 0.093 | 0.073 | 0.058 |
| Autism spectrum disorder | -0.007 | 0.032 | 0.005 | -0.009 | -0.011 | 0.008 | 0.003 | 0.005 | 0.001 | 0.014 | 0.003 | 0.026 |
| Major depressive disorder | 0.022 | 0.045 | 0.009 | -0.003 | 0.031 | 0.020 | 0.022 | 0.009 | 0.034 | 0.029 | 0.021 | 0.028 |
| N | 7927 | 8084 | 7911 | 8030 | 8146 | 7961 | 8026 | 7969 | 8015 | 8027 | 8015 | 8015 |
| Mean | 0.31 | 1.42 | 0.89 | 0.89 | 3.50 | 0.49 | 0.75 | 0.15 | 0.57 | 2.47 | 2.50 | 2.84 |
| (SD) | (1.02) | (1.86) | (2.01) | (1.61) | (2.52) | (1.09) | (1.53) | (0.56) | (1.05) | (3.64) | (3.72) | (3.73) |

**Supplementary Table II.** Descriptive statistics including correlations between variables at age 13 years

| | 1. | 2. | 3. | 4. | 5. | 6. | 7. | 8. | 9. | 10. | 11. | 12. |
|---|---|---|---|---|---|---|---|---|---|---|---|---|
| 1. Depression | 1 | | | | | | | | | | | |
| 2. Generalized anxiety | 0.305 | 1 | | | | | | | | | | |
| 3. Separation Anxiety | 0.275 | 0.338 | 1 | | | | | | | | | |
| 4. Social anxiety | 0.206 | 0.275 | 0.207 | 1 | | | | | | | | |
| 5. Specific phobia | 0.114 | 0.357 | 0.253 | 0.249 | 1 | | | | | | | |
| 6. Irritability | 0.338 | 0.198 | 0.241 | 0.200 | 0.100 | 1 | | | | | | |
| 7. Headstrong | 0.278 | 0.131 | 0.210 | 0.152 | 0.050 | 0.810 | 1 | | | | | |
| 8. Hurtful | 0.234 | 0.113 | 0.190 | 0.145 | 0.061 | 0.621 | 0.642 | 1 | | | | |
| 9. Conduct disorder | 0.256 | 0.092 | 0.124 | 0.108 | 0.016 | 0.440 | 0.502 | 0.453 | 1 | | | |
| 10. Hyperactivity-impulsivity | 0.223 | 0.163 | 0.203 | 0.144 | 0.068 | 0.462 | 0.525 | 0.383 | 0.343 | 1 | | |
| 11. Inattention | 0.274 | 0.170 | 0.174 | 0.230 | 0.089 | 0.471 | 0.516 | 0.345 | 0.396 | 0.664 | 1 | |
| 12. Social-communication | 0.293 | 0.184 | 0.196 | 0.213 | 0.113 | 0.618 | 0.648 | 0.483 | 0.457 | 0.573 | 0.565 | 1 |
| Polygenic risk scores | | | | | | | | | | | | |
| Schizophrenia | 0.016 | -0.004 | 0.031 | 0.025 | -0.006 | 0.031 | 0.042 | -0.004 | 0.018 | 0.020 | 0.070 | 0.051 |
| Attention-deficit/hyperactivity disorder | 0.037 | -0.008 | 0.054 | 0.007 | 0.028 | 0.060 | 0.075 | 0.047 | 0.081 | 0.091 | 0.091 | 0.062 |
| Autism spectrum disorder | 0.022 | 0.012 | 0.027 | 0.013 | -0.001 | -0.002 | 0.016 | -0.005 | 0.004 | 0.004 | 0.022 | -0.002 |
| Major depressive disorder | 0.051 | 0.041 | 0.028 | 0.013 | 0.026 | 0.025 | 0.015 | 0.004 | 0.015 | 0.010 | 0.034 | 0.015 |
| N | 6825 | 6910 | 6340 | 6908 | 6969 | 6870 | 6901 | 6844 | 6922 | 6939 | 6921 | 6932 |
| Mean | 0.42 | 1.86 | 0.49 | 1.25 | 3.70 | 0.47 | 0.61 | 0.12 | 0.60 | 1.45 | 2.65 | 2.56 |
| (SD) | (1.32) | (2.30) | (1.47) | (1.91) | (3.03) | (1.10) | (1.43) | (0.50) | (1.31) | (2.87) | (3.95) | (3.65) |

**Supplementary Table III.** Factor loadings and omega reliability coefficients at age 13 years

| | P | E | B | N |
|---|---|---|---|---|
| Factor loadings | | | | |
| Depression | 0.393 | 0.306 | | |
| Generalized anxiety | 0.213 | 0.664 | | |
| Separation Anxiety | 0.269 | 0.436 | | |
| Social anxiety | 0.241 | 0.365 | | |
| Specific phobia | 0.099 | 0.500 | | |
| Irritability | 0.777 | | 0.424 | |
| Headstrong | 0.824 | | 0.399 | |
| Hurtful | 0.636 | | 0.296 | |
| Conduct disorder | 0.601 | | | |
| Hyperactivity-impulsivity | 0.616 | | | 0.596 |
| Inattention | 0.632 | | | 0.464 |
| Social-communication | 0.784 | | | 0.157 |
| Omega reliability coefficients | | | | |
| Omega ($\omega$) | 0.869 | | | |
| Omega hierarchical ($\omega_H$) | 0.213 | | | |
| Omega subscale ($\omega_S$) | | 0.901 | 0.919 | 0.626 |
| Omega hierarchical subscale ($\omega_{HS}$) | | 0.892 | 0.722 | 0.165 |

P = general psychopathology, E=emotional, B=behavioural, N=neurodevelopmental

**Supplementary Table IV.** Univariable associations between genetic risk and the factor model at age 7 years

| | Schizophrenia PRS | | | | ADHD PRS | | | | ASD PRS | | | | Depression PRS | | | |
|---|---|---|---|---|---|---|---|---|---|---|---|---|---|---|---|---|
| | β | SE | p | $R^2$ | β | SE | p | $R^2$ | β | SE | p | $R^2$ | β | SE | p | $R^2$ |
| General psychopathology | 0.048 | 0.018 | 0.006 | 0.002 | 0.093 | 0.019 | <0.001 | 0.009 | 0.026 | 0.018 | 0.136 | 0.001 | 0.041 | 0.017 | 0.019 | 0.002 |
| Emotional problems | 0.042 | 0.019 | 0.026 | 0.002 | -0.039 | 0.020 | 0.048 | 0.002 | 0.002 | 0.019 | 0.902 | <0.001 | 0.033 | 0.019 | 0.078 | 0.001 |
| Behavioural problems | 0.010 | 0.027 | 0.702 | <0.001 | -0.057 | 0.033 | 0.083 | 0.003 | -0.034 | 0.027 | 0.209 | 0.001 | -0.024 | 0.027 | 0.368 | 0.001 |
| Neurodevelopmental problems | -0.003 | 0.023 | 0.910 | <0.001 | 0.035 | 0.024 | 0.144 | 0.001 | -0.014 | 0.022 | 0.545 | <0.001 | 0.000 | 0.022 | 0.999 | <0.001 |

ADHD=attention-deficit/hyperactivity disorder, ASD=autism spectrum disorder, PRS=polygenic risk score

**Supplementary Table V.** Correlations between factors across ages

| N=5999 | P<br>age 13 | E<br>age 13 | B<br>age 13 | N<br>age 13 |
|---|---|---|---|---|
| General psychopathology (P), age 7 | 0.762 | -0.055 | -0.444 | -0.039 |
| Emotional problems (E), age 7 | -0.057 | 0.828 | 0.080 | 0.046 |
| Behavioural problems (B), age 7 | -0.305 | 0.120 | 0.871 | 0.347 |
| Neurodevelopmental problems (N), age 7 | -0.115 | 0.050 | 0.328 | 0.907 |

**Supplementary Table VI.** Correlations between polygenic risk scores

| N=6166 | Schizophrenia | ADHD | ASD | Depression |
|---|---|---|---|---|
| Schizophrenia | 1 | | | |
| ADHD | 0.040 | 1 | | |
| ASD | 0.052 | 0.214 | 1 | |
| Depression | 0.086 | 0.128 | 0.109 | 1 |

**Supplementary Table VII.** Differences between those with and without polygenic risk score data on variables included in the inverse probability weighting model

| | With PRS data (N=7495) | Without PRS data (N=6298) | Association with missingness |
|---|---|---|---|
| Child gender, female | 48.317% | 48.472% | OR=0.99 (0.93-1.06), p=0.856 |
| Child birth weight | Mean=3437.326 (SD=532.572) | Mean=3361.634 (SD=568.202) | OR=1.00 (1.00-1.00)*, p<0.001 |
| Maternal age | Mean=28.700 (SD=4.755) | Mean=27.119 (SD=5.086) | OR=0.94 (0.93-0.94), p<0.001 |

PRS=polygenic risk score. N=13793 for maternal age and child gender, N=13619 for child birth weight. *OR=0.9997 (0.9997-0.9998)

**Supplementary Table VIII.** Multivariable associations between genetic risk and the factor model at age 7 years using inverse probability weighting

| | Schizophrenia PRS | | | ADHD PRS | | | ASD PRS | | | MDD PRS | | | |
|---|---|---|---|---|---|---|---|---|---|---|---|---|---|
| | β | SE | p | β | SE | p | β | SE | p | β | SE | p | $R^2$ |
| General psychopathology | 0.042 | 0.018 | 0.019 | 0.092 | 0.020 | <0.001 | -0.001 | 0.019 | 0.943 | 0.027 | 0.018 | 0.126 | 0.012 |
| Emotional problems | 0.041 | 0.020 | 0.036 | -0.045 | 0.021 | 0.031 | 0.007 | 0.019 | 0.729 | 0.033 | 0.020 | 0.094 | 0.005 |
| Behavioural problems | 0.025 | 0.029 | 0.388 | -0.053 | 0.037 | 0.147 | -0.024 | 0.030 | 0.422 | -0.014 | 0.029 | 0.618 | 0.005 |
| Neurodevelopmental problems | 0.000 | 0.024 | 0.995 | 0.040 | 0.026 | 0.131 | -0.019 | 0.024 | 0.421 | -0.001 | 0.022 | 0.961 | 0.002 |

ASD=autism spectrum disorder, ADHD=attention-deficit/hyperactivity disorder, MDD=major depressive disorder, PRS=polygenic risk score

Wray NR, Ripke S, Mattheisen M, Trzaskowski M, Byrne EM, Abdellaoui A, Adams MJ, Agerbo E, Air TM, Andlauer TMF, Bacanu SA, Baekvad-Hansen M, Beekman AFT, Bigdeli TB, Binder EB, Blackwood DRH, Bryois J, Buttenschon HN, Bybjerg-Grauholm J, Cai N, Castelao E, Christensen JH, Clarke TK, Coleman JIR, Colodro-Conde L, Couvy-Duchesne B, Craddock N, Crawford GE, Crowley CA, Dashti HS, Davies G, Deary IJ, Degenhardt F, Derks EM, Direk N, Dolan CV, Dunn EC, Eley TC, Eriksson N, Escott-Price V, Kiadeh FHF, Finucane HK, Forstner AJ, Frank J, Gaspar HA, Gill M, Giusti-Rodriguez P, Goes FS, Gordon SD, Grove J, Hall LS, Hannon E, Hansen CS, Hansen TF, Herms S, Hickie IB, Hoffmann P, Homuth G, Horn C, Hottenga JJ, Hougaard DM, Hu M, Hyde CL, Ising M, Jansen R, Jin F, Jorgenson E, Knowles JA, Kohane IS, Kraft J, Kretzschmar WW, Krogh J, Kutalik Z, Lane JM, Li Y, Li Y, Lind PA, Liu X, Lu L, MacIntyre DJ, MacKinnon DF, Maier RM, Maier W, Marchini J, Mbarek H, McGrath P, McGuffin P, Medland SE, Mehta D, Middeldorp CM, Mihailov E, Milaneschi Y, Milani L, Mill J, Mondimore FM, Montgomery GW, Mostafavi S, Mullins N, Nauck M, Ng B, Nivard MG, Nyholt DR, O'Reilly PF, Oskarsson H, Owen MJ, Painter JN, Pedersen CB, Pedersen MG, Peterson RE, Pettersson E, Peyrot WJ, Pistis G, Posthuma D, Purcell SM, Quiroz JA, Qvist P, Rice JP, Riley BP, Rivera M, Saeed Mirza S, Saxena R, Schoevers R, Schulte EC, Shen L, Shi J, Shyn SI, Sigurdsson E, Sinnamon GBC, Smit JH, Smith DJ, Stefansson H, Steinberg S, Stockmeier CA, Streit F, Strohmaier J, Tansey KE, Teismann H, Teumer A, Thompson W, Thomson PA, Thorgeirsson TE, Tian C, Traylor M, Treutlein J, Trubetskoy V, Uitterlinden AG, Umbricht D, Van der Auwera S, van Hemert AM, Viktorin A, Visscher PM, Wang Y, Webb BT, Weinsheimer SM, Wellmann J, Willemsen G, Witt SH, Wu Y, Xi HS, Yang J, Zhang F, eQtlgen, andMe, Arolt V, Baune BT, Berger K, Boomsma DI, Cichon S, Dannlowski U, de Geus ECJ, DePaulo JR, Domenici E, Domschke K, Esko T, Grabe HJ, Hamilton SP, Hayward C, Heath AC, Hinds DA, Kendler KS, Kloiber S, Lewis G, Li QS, Lucae S, Madden PFA, Magnusson PK, Martin NG, McIntosh AM, Metspalu A, Mors O, Mortensen PB, Muller-Myhsok B, Nordentoft M, Nothen MM, O'Donovan MC, Paciga SA, Pedersen NL, Penninx B, Perlis RH, Porteous DJ, Potash JB, Preisig M, Rietschel M, Schaefer C, Schulze TG, Smoller JW, Stefansson K, Tiemeier H, Uher R, Volzke H, Weissman MM, Werge T, Winslow AR, Lewis CM, Levinson DF, Breen G, Borglum AD, Sullivan PF, Major Depressive Disorder Working Group of the Psychiatric Genomics C (2018) Genome-wide association analyses identify 44 risk variants and refine the genetic architecture of major depression. Nat Genet 50(5):668-681

Euesden J, Lewis CM, O'Reilly PF (2015) PRSice: Polygenic Risk Score software. Bioinformatics 31(9):1466-1468

Seaman SR, White IR, Copas AJ, Li L (2012) Combining multiple imputation and inverse-probability weighting. Biometrics 68(1):129-137
